# Cardiomyocyte-specific ACSL1 Deficiency Prevents Cardiac Lipotoxicity and Alleviates Heart Dysfunction in the *ob/ob* Model of Obesity

**DOI:** 10.1101/2020.01.24.918961

**Authors:** Florencia Pascual, Trisha J. Grevengoed, Liyang Zhao, Monte S. Willis, Rosalind A. Coleman

## Abstract

Cardiac lipotoxicity is associated with structural remodeling and functional changes that are features of obesity-related cardiomyopathy. Both high fat diet and the *ob/ob* mutation lead to increased fatty acid (FA) uptake, elevated triacylglycerol (TAG) content, hypertrophy, and systolic and diastolic dysfunction in murine hearts. Cardiomyocyte-specific long-chain acyl-CoA synthetase 1 (ACSL1) deficiency (*Acsl1*^*H*-/-^) results in a 90% reduction in FA activation, suggesting that *Acsl1* ablation might alleviate obesity-associated myocardial dysfunction. Double knockout *ob-Acsl1*^*H*-/-^ and *ob-Acsl1*^*flox/flox*^ control mice were treated with tamoxifen at 20 weeks of age; heart function, TAG content, and relevant gene expression were assessed immediately before and 2 and 5 weeks after treatment. Heart weights initially increased in lean and obese *Acsl1*^*H*-/-^ mice, but normalized in *ob-Acsl1*^*H*-/-^ mice by 5 weeks. Ventricular TAG content was decreased by 51% and 61% in *ob-Acsl1*^*H*-/-^ mice 2 and 5 weeks after *Acsl1* knockout induction, respectively. Moreover, ACSL1 knockout resulted in increased survival of *ob/ob* mice, suggesting that lack of ACSL1 protected obese hearts subjected to stress. Our results indicate that partial knockdown of cardiac ACSL1 is sufficient to reverse cardiac TAG accumulation and to ameliorate heart dysfunction even in the context of established obesity-related cardiomyopathy.

## Introduction

The adult healthy heart has a marked preference for oxidation of long-chain fatty acids (FA) to fuel its continuous contractile activity, but can readily switch to alternative energy substrates to preserve cardiac output under varying physiological conditions [1]. In contrast, pathologies such as metabolic syndrome, obesity and diabetes are characterized by a loss of cardiac metabolic flexibility as well as alterations in FA metabolism [1-3] that lead to excess cardiac lipid storage [2-5]. Experimental and clinical studies have implicated myocardial lipid accumulation in the development of obesity-associated cardiomyopathies [1]; however, the contribution of specific lipid species to disease pathogenesis has yet to be fully elucidated. Whereas early studies linked increased TAG content to cardiac lipotoxicity, recent work has focused on the effect of diacylglycerol (DAG) and ceramide, which can sometimes (but not always) accompany increased intramyocardial TAG.

Excess cardiac TAG accumulation has been associated with altered lipid signaling, accumulation of reactive oxygen species, ER stress, and impaired mitochondrial function [1]. Moreover, TAG-associated cardiac lipotoxicity in humans correlates with structural remodeling and functional changes that are features of obesity-related cardiomyopathy, such as left ventricular hypertrophy and impaired myocardial contractility [1]. For instance, impaired TAG lipolysis in a mouse model of *Atgl* deficiency results in severe cardiac lipid accumulation, cardiac dysfunction, and premature death, in part due to a defect in PPARα activation [6,7]. In rodent models, both high fat diet [8,9] and leptin deficiency [10,11] result in increased cardiac FA uptake, elevated TAG content, cardiac hypertrophy, and systolic and diastolic dysfunction. Conversely, obesity-associated heart dysfunction is ameliorated by reducing cardiac FA uptake [12,13], and by increasing FA oxidation [14], lipoprotein secretion [15], or esterification of free FA [16].

ACSLs catalyze the conversion of FA to acyl-CoAs, an obligatory step for their downstream use as either energy substrates in β-oxidation or as building blocks in the synthesis of TAG and other complex lipids. ACSL1 is the major cardiac ACSL isoform in a family of 5 closely-related isoenzymes, contributing 50-90% of the total ACS activity. Cardiomyocyte-specific *Acsl1* ablation (*Acsl1*^*H-/-*^) leads to > 90% reduction in mitochondrial FA oxidation, a 35% decrease in lipid uptake, and an 8-fold increase in glucose uptake and use [17,18]. Notably, *Acsl1* knockout does not affect cardiac TAG levels, suggesting that ACSL1-mediated activation channels FA specifically towards oxidation in the heart. However, long-term ACSL1 inactivation is also accompanied by deleterious effects, including glucose intolerance, development of mTORC1-mediated pathological hypertrophy, and diastolic dysfunction [17,18]. In contrast, partial (∼60%) cardiac ACSL1 knockdown is sufficient to elicit cardiac-specific metabolic derangements and the development of hypertrophy without affecting cardiac function [19]. We therefore hypothesized that partial *Acsl1* ablation could reverse ventricular TAG accumulation and ameliorate myocardial dysfunction in the *ob/ob* model of cardiomyopathy. Our findings suggest that preventing or reversing cardiac lipid accumulation in the context of established lipotoxicity contributes to improved cardiac function, revealing *Acsl1* as a potential target in the development of cardiovascular therapies for obesity-associated cardiomyopathies.

## Materials and Methods

### Animal Treatment

The University of North Carolina Institutional Animal Care and Use Committee approved all protocols. Mice were housed in a pathogen-free barrier facility (12-h light/12-h dark cycle) with free access to water and food (Prolab RMH 3000 SP76 chow). *Acsl1*^*H-/-*^ [17] and *ob*/+ [20] mice were bred to generate an *ob-Acsl1*^*H-/-*^ double knockout model (Fig. 1A). At 20 weeks of age, *ob-Acsl1*^*H-/-*^ and lean and obese *Acsl1*^*flox/flox*^ littermate control mice were injected intraperitoneally with tamoxifen (3 mg/40 g of body weight) (Sigma) dissolved in corn oil (20 mg/ml) for 4 consecutive days; all mice were injected with tamoxifen to adequately control for both the intrinsic effects of Cre/tamoxifen [21] and the TAG species alterations that result from corn oil vehicle treatment. Systolic and diastolic heart function was examined by M-mode echocardiography and mitral Doppler analysis immediately before, and 2 and 5 weeks after tamoxifen treatment. Two and five weeks after tamoxifen induction, animals were anesthetized with 2,2,2-tribromoethanol (Avertin) and tissues were removed and snap-frozen in liquid nitrogen (Fig. 1B). To isolate total membranes, left ventricles were homogenized with 10 up-and-down strokes using a motor-driven Teflon pestle and glass mortar in ice-cold medium I (MedI) buffer (10 mM Tris [pH 7.5], 1 mM EDTA, 250 mM sucrose, 1 mM dithiothreitol [DTT], plus protease and phosphatase inhibitor cocktail [Roche]). Homogenates were centrifuged at 100,000 ×*g* for 1 h at 4 °C; the membrane pellet was then resuspended in MedI buffer. Protein content was determined by the bicinchoninic acid (BCA) assay (Pierce) with bovine serum albumin (BSA) as the standard. Plasma was collected from mice in 5% 0.5 M EDTA. Plasma TAG was measured with a colorimetric assay (Sigma).

**Figure 1.**
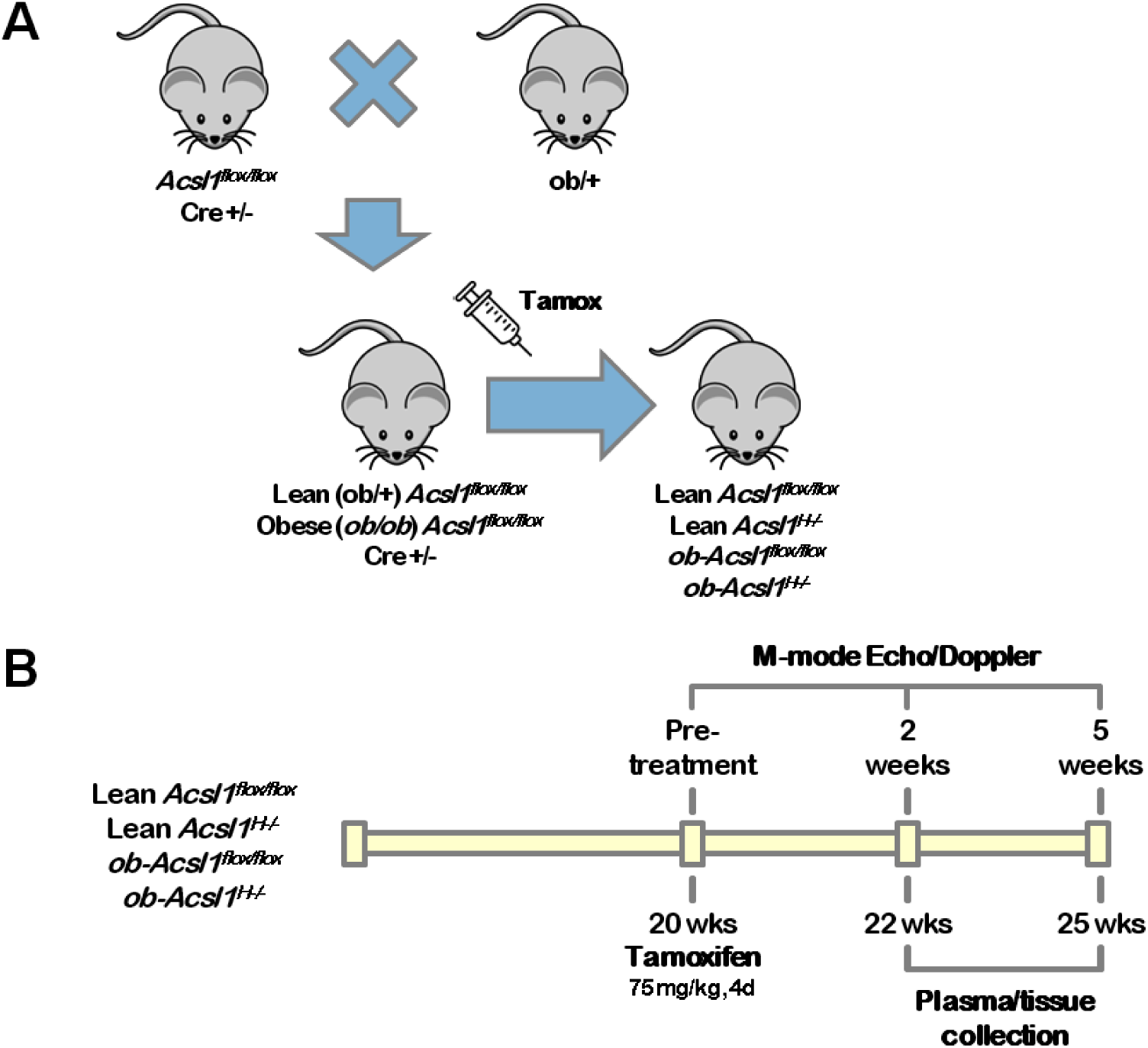
Study design. *A*, Mice with *Loxp* sequences inserted on either side of exon 2 in the *Acsl1* gene were interbred with mice in which Cre expression is driven by a α-myosin heavy chain promoter induced by tamoxifen to generate tamoxifen-inducible heart-specific (*Acsl1*^*H-/-*^) *Acsl1* knockout mice [33]. To obtain leptin deficiency (*ob/ob*) on a background of tamoxifen-inducible *Acsl1* knockout (*ob-Acsl1*^*H-/-*^), *Acsl1*^*H-/-*^ and ob/+ mice were crossed to obtain floxed mice that had either lean (ob/+) or obese (*ob/ob*) phenotypes. *Acsl1* deficiency was induced by tamoxifen (*tamox*) injection. All offspring were genotyped by PCR techniques. *B*, 20 week-old mice obtained through the breeding scheme described above were injected with tamoxifen for 4 consecutive days. Systolic and diastolic heart function was examined by M-mode echocardiography and mitral Doppler analysis immediately before, and 2 and 5 weeks after tamoxifen treatment; plasma and tissues were collected two and five weeks after *Acsl1* ablation.

### Echocardiography and Doppler Analysis

Cardiac echocardiography was performed (blinded to mouse type and treatment) on unanesthetized mice with the VisualSonics Vevo 770 or Vevo 2100 ultrasound biomicroscopy system as described [17,18]. M-mode images of the left ventricle were analyzed using VisualSonics software. Doppler analysis of the mitral valve to determine the mitral inflow velocity was performed on lightly anesthetized mice (2% (vol/vol) isoflurane/100% oxygen) as described [18,22]. Mitral valve flow Doppler was acquired by positioning the transducer in an epigastric position angled cranially in a supine mouse at 45° to achieve an apical four-chamber view. Peak E and A heights were determined on mitral valve sequential waveforms in at least 5 waveforms using VisualSonics software. The mean performance index (MPI) was calculated as the isovolumetric contraction time (ICT) plus the isovolumetric relaxation time (IRT) divided by the ejection time ((ICT + IRT)/ET) to determine if either systolic or diastolic dysfunction was present [22,23]. E-wave deceleration time (DT) was also measured. Doppler measurement data represent 3–6 averaged cardiac cycles from at least 3 scans per mouse.

### Acsl Assay

ACSL initial rates were measured with 50 μM [1-^14^C]palmitic acid (Perkin Elmer), 10 mM ATP, and 0.25 mM coenzyme A (CoA) in total membrane fractions from ventricles (2–6 μg protein) as described [17]. No ACSL activity was measurable in the soluble (cytosolic) fraction.

### RT-PCR

Total RNA was isolated from left ventricles (RNeasy Fibrous Tissues Kit, Qiagen) and cDNA was synthesized (Applied Biosystems high-capacity cDNA reverse transcription kit). cDNA was amplified by real-time PCR using SYBR Green (Applied Biosystems) detection with specific primers for the gene of interest. Results were normalized to the housekeeping gene *Gapdh* and expressed as arbitrary units of 2^-ΔΔCT^ relative to the control group (Table 1).

**Table 1.**
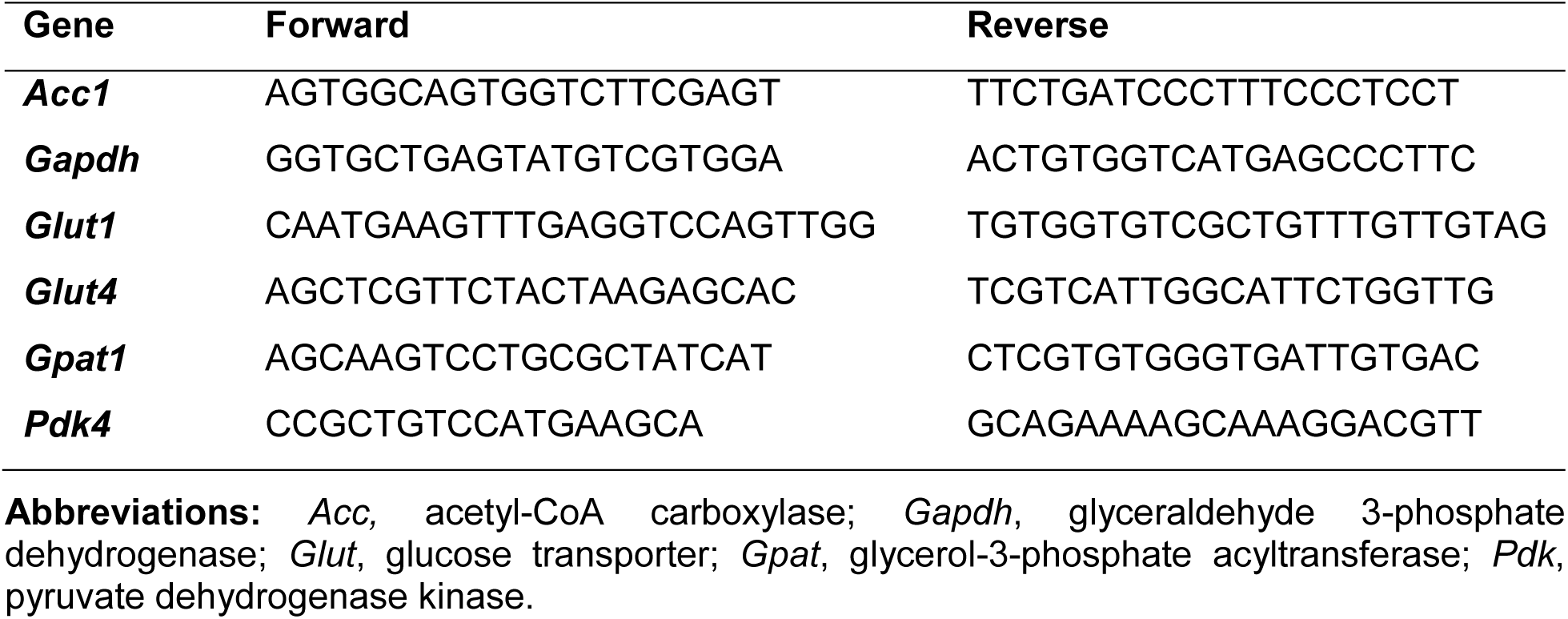
Quantitative RT-PCR primers.

### Immunoblots

Total protein lysates from left ventricles were isolated in lysis buffer (20 mM Tris base, 1% Triton X-100, 50 mM NaCl, 250 mM sucrose, 50 mM NaF, 5 mM Na2P2O7, plus Halt™ protease and phosphatase inhibitor cocktail [Thermo Scientific]) or MedI buffer. Equal amounts of protein (100 μg) were loaded and resolved on 10% SDS polyacrylamide gels and transferred to nitrocellulose membranes. Blots were probed with antibodies against ACSL1 (catalog no. 4047), phosphorylated p70 (P-p70) S6 kinase (S6K) (catalog no. 9234), and phosphorylated 4E-BP1 (catalog no. 2855), and were then stripped and reprobed with p70 S6K (catalog no. 2708) or 4E-BP1 (catalog no. 9644) antibodies, respectively (all antibodies from Cell Signaling,). GAPDH (Abcam, catalog no. ab8245) was used as loading control.

### Histology

Heart ventricles were fixed for 24h in PBS containing 4% paraformaldehyde, transferred to 70% ethanol, embedded in paraffin, serial sectioned, and stained with Masson’s trichrome. Slides were scanned using an Aperio ScanScope (Aperio Technologies); images of left ventricular myocardium were selected for figure 2F.

**Figure 2.**
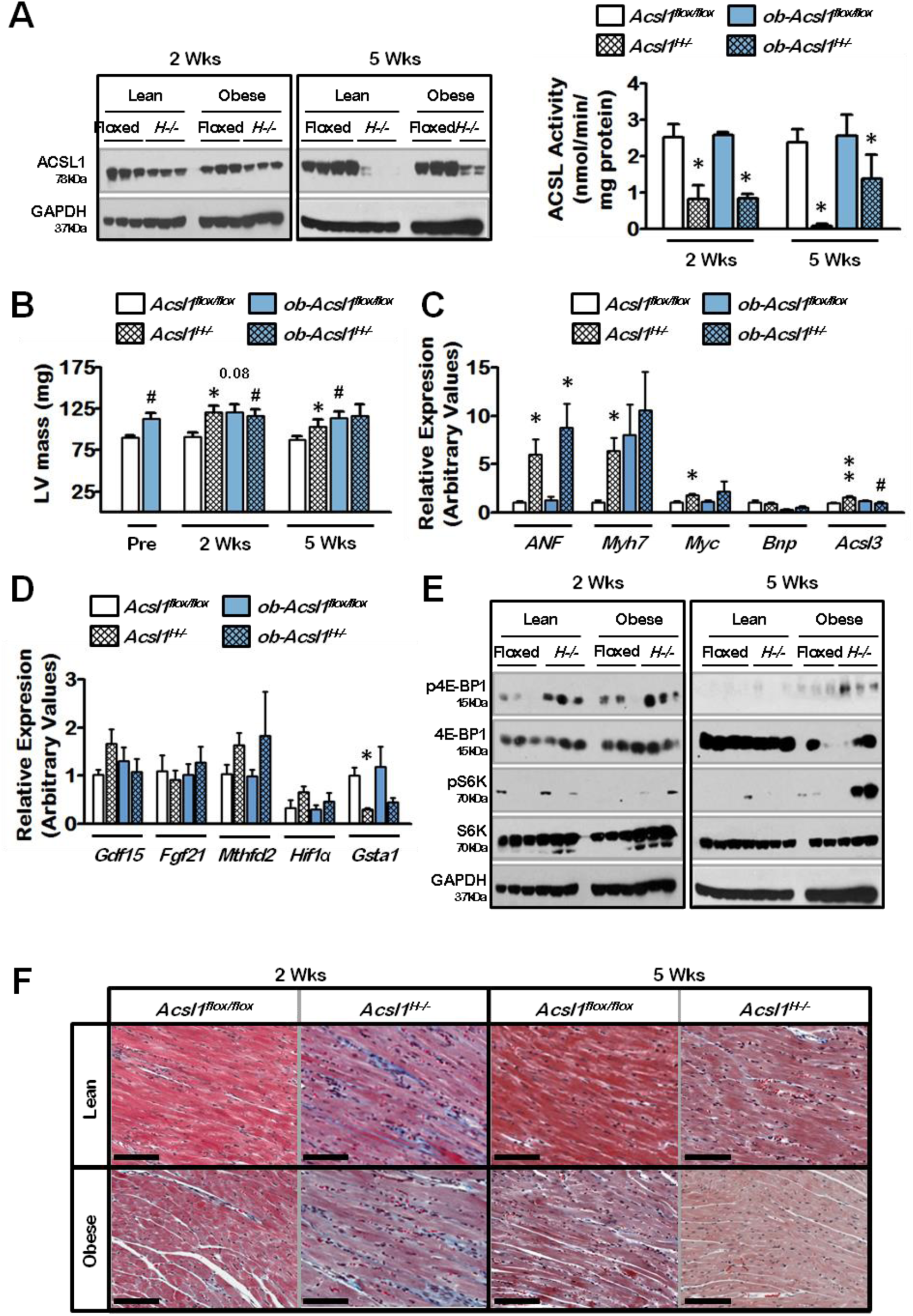
*Acsl1* abrogation does not significantly exacerbate mTOR-mediated cardiac hypertrophy or fibrosis in the context of obesity-associated cardiomyopathy. *A*, Ventricular ACSL1 protein levels and total ACSL activity 2 and 5 weeks after tamoxifen treatment. n = 3-12. *B*, Hypertrophy as determined by transthoracic echocardiographic assessment of left ventricle (*LV*) mass in obese and lean floxed and *Acsl1*^*H-*/-^ (*H*^*-/-*^) knockout mice immediately before (*Pre*), 2 and 5 weeks after tamoxifen treatment. n= 3-9. mRNA abundance of fetal gene markers (*C*) and mTOR-responsive genes (*D*) in ventricles from control and *ob-Acsl1*^*H*−/−^ mice 2 weeks after knockout induction. n = 5. *E*, Representative immunoblots from *ob-Acsl1*^*H-*/-^ and control mice 2 and 5 weeks after tamoxifen treatment. Blots were probed for p-4EBP1 and p-S6K, known downstream targets of activated mTOR. *F*, Masson’s trichrome staining of left ventricular myocardium from lean and obese mice 2 and 5 weeks after cardiac-specific *Acsl1* knockdown; original magnification 20×, scale bar = 100 μm, n = 2. The values reported are means ± SEM. *, p ≤ 0.05 and **, p ≤ 0.01 compared with floxed controls; #, p ≤ 0.05 compared with lean phenotype controls.

### Statistics

Values are expressed as mean ± standard error of the mean (SEM). Differences between genotypes were evaluated by two-tailed unpaired Student’s t-test. Differences between lean and obese groups were evaluated by two-way ANOVA with Tukey multiple-comparison post-tests. All statistical analyses were performed using GraphPad Prism (version 6.0). For all tests, *p* < 0.05 was considered significant.

## Results

### Acsl1 ablation does not exacerbate obesity-induced cardiac hypertrophy or fibrosis

Disruption of leptin signaling has been linked to cardiac hypertrophy [5,24,25]. Total cardiac-specific *Acsl1* ablation is achieved 10 weeks after tamoxifen treatment and results in >90% reduction in cardiac ACSL activity and FA oxidation, a compensatory shift towards glucose uptake and use, and the development of mTOR-responsive hypertrophy and diastolic dysfunction [17,18]. Conversely, partial deficiency of ACSL1-mediated FA activation observed 2 weeks after tamoxifen treatment induces changes in cardiac metabolism and size without affecting heart function [19]. Thus, we set out to examine whether partial *Acsl1* inactivation at both early (2 weeks) and intermediate (5 weeks) time points could ameliorate heart dysfunction in the context of obesity-associated cardiomyopathy. We induced cardiac-specific *Acsl1* knockdown in 20-week old obese mice and in lean littermate controls, and examined hearts 2 and 5 weeks after tamoxifen treatment (Fig. 2). ACSL1 protein and ACSL activity were still detected 2 weeks after *Acsl1* inactivation, but these levels were drastically reduced at 5 weeks (Fig. 2A). Ventricular ACSL activity was reduced by ∼60% 2 weeks after tamoxifen treatment, whereas <10% of normal activity was observed in hearts 5 weeks after knockout induction, similar to the total *Acsl1* ablation that occurs 10 weeks after tamoxifen treatment [17,18]. As observed in numerous earlier studies [4,5,24-26], leptin deficiency was accompanied by left ventricular hypertrophy (Fig. 2B, “*Pre*”), with left ventricle masses that were 25% greater in 20-wk old *ob/ob* mice compared to lean littermate controls. *Acsl1* abrogation resulted in enlarged hearts in lean mice 2 and 5 weeks after tamoxifen treatment, but did not contribute to increased hypertrophy in obese mice. In addition, partial ACSL1 knockdown did not alter the expression of pathological hypertrophy markers (Fig. 2C) or mTOR targets (Fig. 2D), while mTOR-mediated phosphorylation of p70 S6 kinase (S6K) was dramatically upregulated in the context of obesity (Fig. 2E). As shown previously [34], short-term inactivation of *Acsl1* resulted in a transient increase in cardiac fibrosis in lean and obese mice 2 weeks after tamoxifen treatment, but did not exacerbate the mild fibrosis observed in hearts from obese mice (Fig. 2F).

### Acsl1 inactivation reduces ventricular stores of TAG without altering systemic lipid metabolism

The cardiac-specific knockdown of *Acsl1* did not affect total body or relative tissue weight in *ob/ob* mice (Fig. 3A). Expression of cardiac FA transporters and *de novo* FA synthesis genes is elevated in *ob/ob* hearts [26], and studies have suggested that increased levels of SREBP-1c and its targets, *Gpat1* and *Acc1*, lead to elevated rates of hepatic FA and TAG synthesis [27]. However, in agreement with a previous study [5], we did not observe differences in cardiac expression of SREBP-1c targets between *ob/ob* and lean controls (Fig. 3B). Moreover, cardiac expression of key genes involved in glucose metabolism and circulating levels of TAG (Fig. 3C) were unaltered, suggesting that partial ablation of cardiac ACSL1 does not affect cardiac or systemic glucose homeostasis and FA synthesis. In contrast, ventricular TAG content was decreased by 51% and 61% in *ob-Acsl1*^*H-*/-^ mice 2 and 5 weeks after *Acsl1* inactivation, respectively (Fig. 3D), indicating that partial cardiac-specific *Acsl1* abrogation is sufficient to reverse established cardiac lipid accumulation.

**Figure 3.**
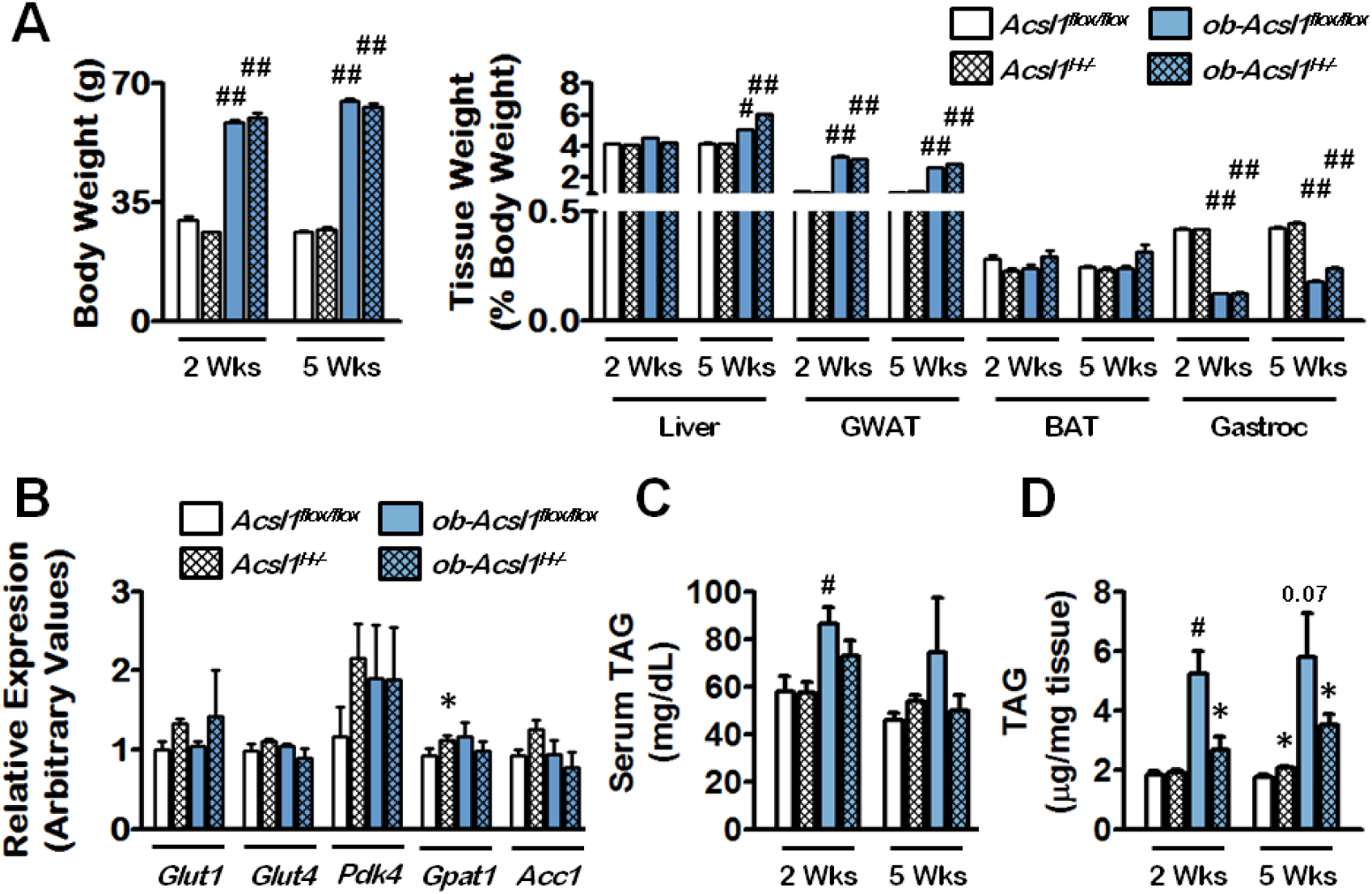
Partial ablation of cardiac *Acsl1* reduces ventricular TAG stores but does not affect systemic lipid metabolism. *A*, Body weight and relative tissue weights of *ob-Acsl1*^*H-*/-^ and control mice 2 and 5 weeks after tamoxifen treatment. n = 4-15. *B*, Expression of key metabolic genes 2 weeks after induction of ACSL1 deficiency. n = 3-7. *C*, TAG content in serum or *D*, ventricles from *ob-Acsl1*^*H-*/-^ and control mice 2 and 5 weeks after tamoxifen treatment. n = 3-12. The values reported are means ± SEM. *, p ≤ 0.05 and **, p ≤ 0.01 compared with littermate floxed controls; #, p ≤ 0.05 and *##*, p ≤ 0.01 compared with lean phenotype controls.

### Acsl1 knockdown ameliorates obesity-induced systolic dysfunction and may be protective in ob/ob hearts subjected to stress

We hypothesized that reversing TAG accumulation could ameliorate obesity-associated cardiomyopathy in *ob/ob* hearts. Ejection Fraction (*EF*), a measure of heart contractility, and Fractional Shortening (*FS*), a parameter of cardiac output, were analyzed by M-mode echocardiography in conscious mice before and 2 and 5 weeks after tamoxifen treatment (Fig. 4A). As expected, cardiac systolic function was decreased in 20-wk old *ob/ob* mice compared to littermate controls. Interestingly, *Acsl1* knockdown resulted in a previously undescribed, transient decline in systolic function 2 weeks after tamoxifen treatment, and contributed slightly to decreased function in *ob/ob* hearts; however, normal systolic function was restored 5 weeks after tamoxifen injection, when cardiac ACSL1 activity was virtually absent and ventricular TAG stores were markedly diminished in obese hearts. Moreover, *ob/ob* hearts exhibited hallmarks of dilated cardiomyopathy and reduced contractility including increased LV diameter, LV volume, and stroke volume, all of which resolved 5 weeks after *Acsl1* knockdown (Table 2). We posit that the temporary dip in function resulted from an initial compensatory process during which cardiac efficiency is decreased through oxygen use for non-contractile purposes [19,28], and that it had not been previously observed due to differences in age and gender of the animals studied [17,18,29]. Diastolic function was evaluated by Doppler analysis of blood flow through the mitral valve on anesthetized lean and obese mice immediately before and 2 and 5 weeks after induction of *Acsl1* knockdown. E/A ratio (*E/A*), a measure of ventricular filling velocity, and Mean Performance Index (*MPI*) were calculated for the assessment of diastolic dysfunction and combined systolic/diastolic performance, respectively (Fig. 4B, C). Obese mice had decreased E/A ratios, indicating mild diastolic dysfunction. Although no significant decreases in diastolic function were observed in obese hearts 2 and 5 weeks after tamoxifen treatment compared with obese hearts at baseline, E/A ratios were decreased in both lean and obese mice at 5 weeks compared to lean hearts at baseline, suggestive of a more general age effect. Obese hearts exhibited decreased E-waves and increased A-waves (Table 3), characteristic of a pseudo-normal filling pattern in which dilation and reduced contractility are compensated by increased left atrial pressure. A trend towards increased MPI in obese mice also suggested impaired diastolic/systolic function that disappeared upon *Acsl1* knockdown. In addition, 44% of *ob/ob* mice subjected to M-mode echocardiography and/or Doppler imaging died during or shortly after these analyses, whereas all *ob-Acsl1*^*H-*/-^ double knockouts survived, indicating that ACSL1 deficiency likely protects obese hearts that are subjected to stress. Given the observed variability in ventricular TAG levels in obese mice (Fig. 3D), we hypothesize that the apparent normal diastolic function we observed in obese mice at weeks 2 and 5 was due to “selection” for those mice whose lower cardiac TAG content made them inherently more resilient to stress.

**Table 2.**
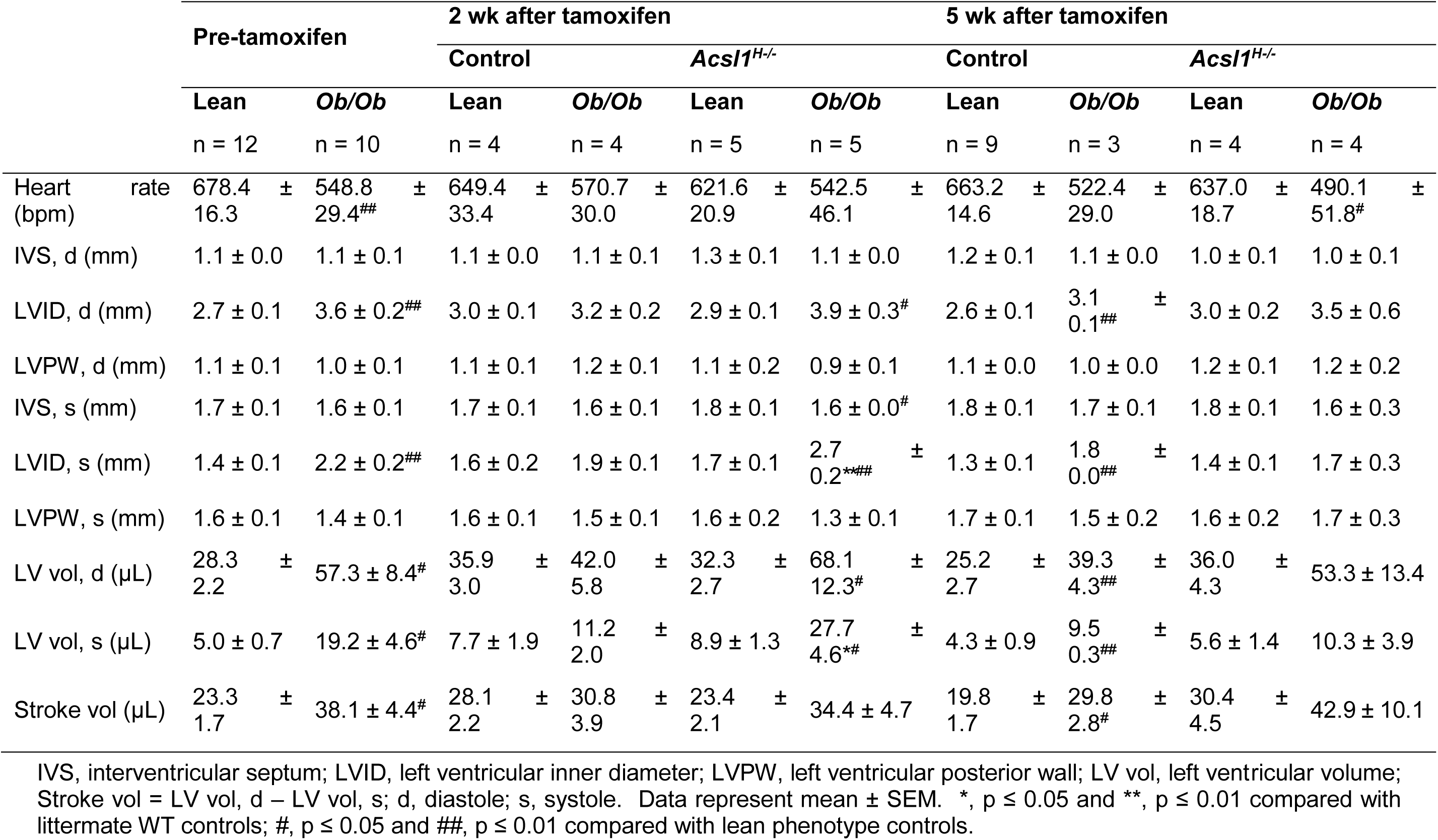
Conscious M-mode echocardiographic analysis of control and *Acsl1*^*H*−/−^ hearts 2 and 5 weeks after tamoxifen induction

**Table 3.**
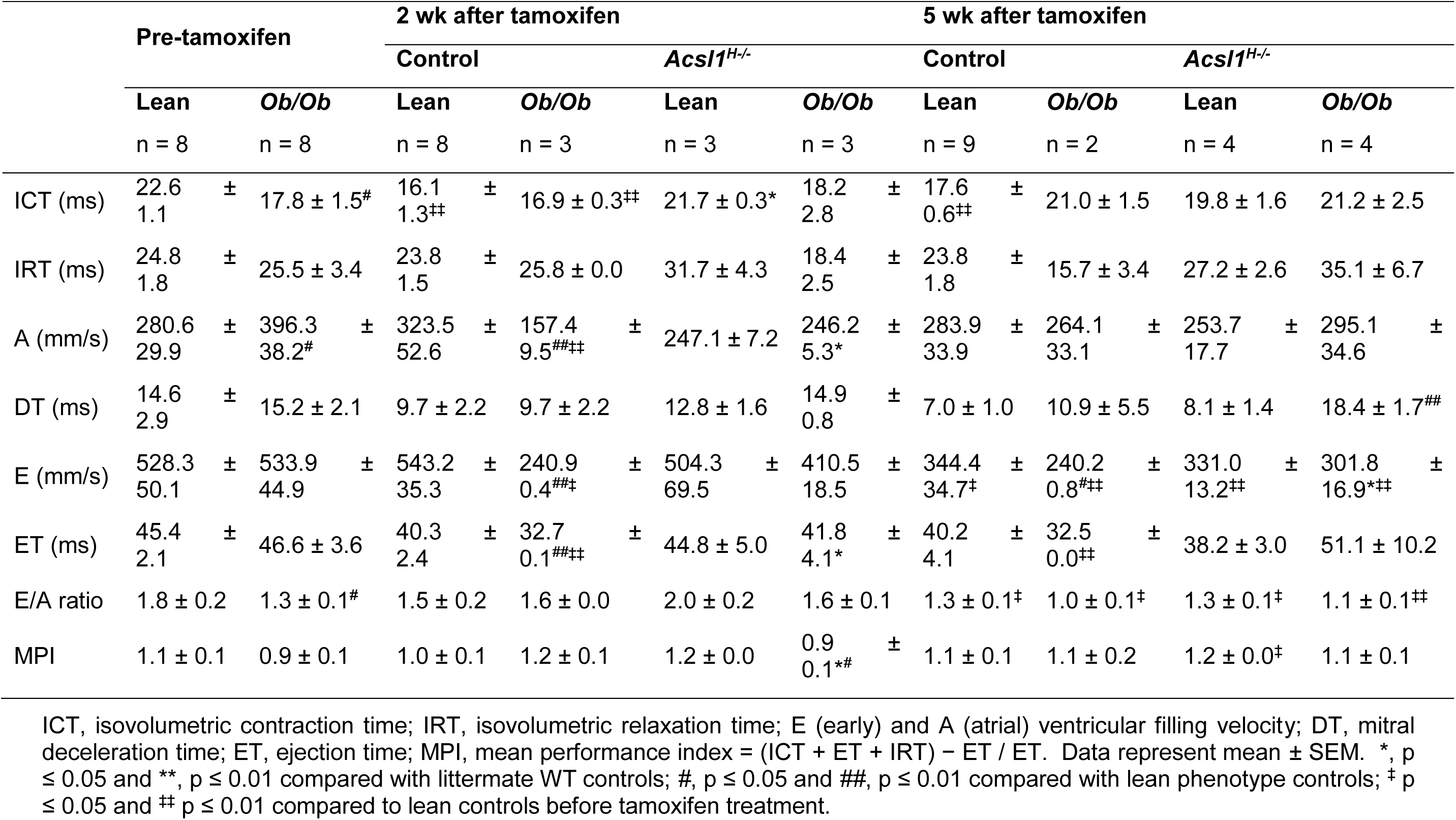
Doppler echocardiographic analysis of control and *Acsl1*^*H*−/−^ hearts 2 and 5 weeks after tamoxifen induction

**Figure 4.**
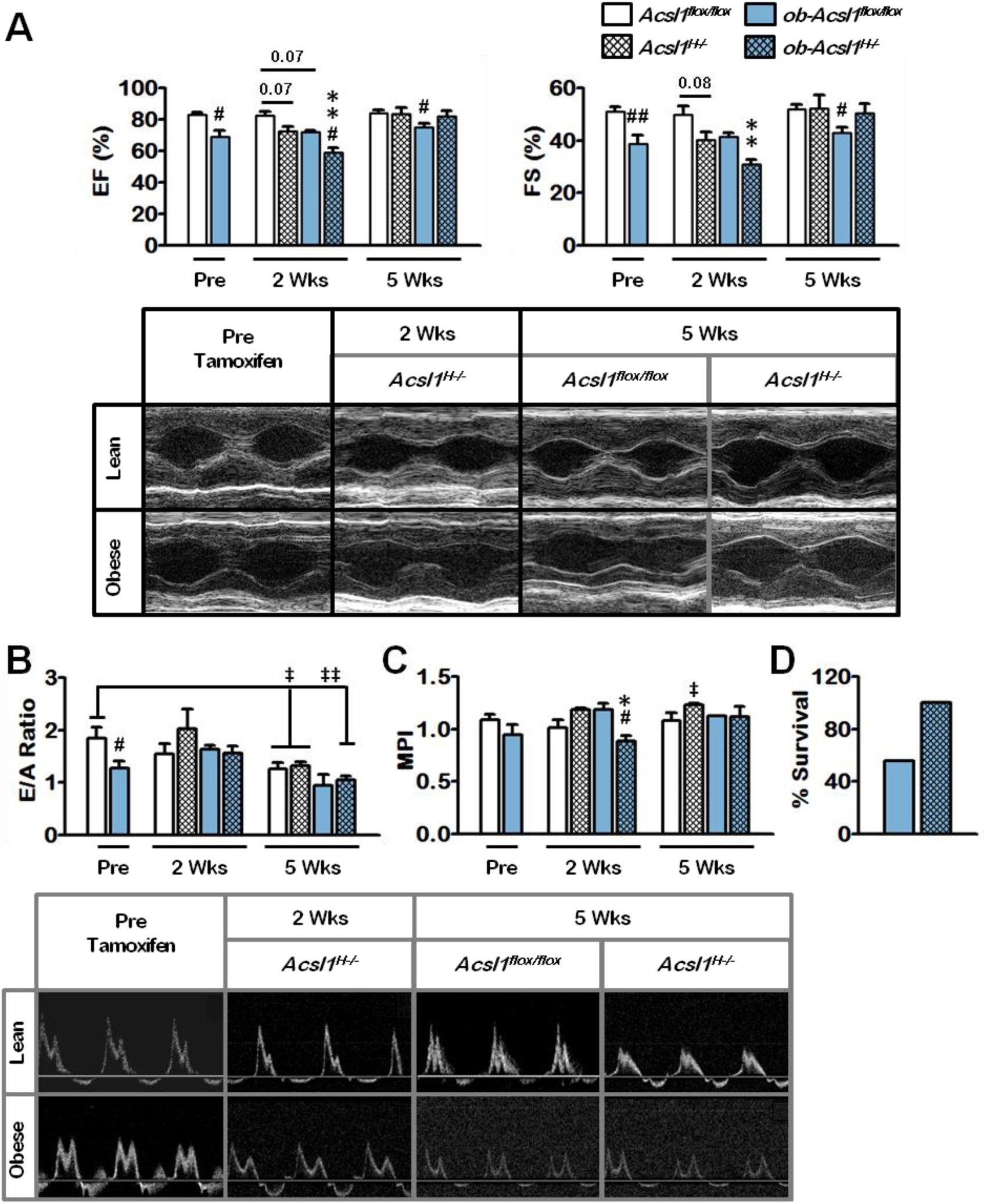
*Acsl1* knockdown rescues systolic dysfunction in the *ob/ob* model of cardiomyopathy and may protect *ob/ob* hearts against echocardiography-induced stress. *A*, Ejection Fraction (*EF*) and Fractional Shortening (*FS*), measures of cardiac systolic function, were analyzed by M-mode echocardiography in conscious *ob-Acsl1*^*H-*/-^ and control mice 2 and 5 weeks after tamoxifen treatment. n = 4-12. *B*, E/A ratio (*E/A*, ratio between early (*E*) and late (atrial-*A*) and *C*, Mean Performance Index (*MPI*) values were calculated from mitral Doppler echocardiography in anesthetized *ob-Acsl1*^*H-*/-^ and control mice 2 and 5 weeks after tamoxifen treatment. n = 2-8. Values reported are means ± SEM. *, p ≤ 0.05 and **, p ≤ 0.01 compared with littermate WT controls; *#*, p ≤ 0.05 and *##*, p ≤ 0.01 compared with lean phenotype controls; ^‡^ p ≤ 0.05 and ^‡‡^ p ≤ 0.01 compared to lean controls before tamoxifen treatment. *D*, Survival analysis of *ob/ob* vs *ob-Acsl1*^*H-/-*^ mice that were subjected to M-mode echocardiography and/or Doppler imaging. 100% = 9.

## Discussion

The ability to readily switch between metabolic substrates confers on cardiac muscle the flexibility required to sustain cardiac output when faced with changes in substrate availability, hormonal status, or physiological conditions. Although the heart can metabolize glucose, lactate, and ketone bodies to produce energy, 60-90% of the cardiac energy requirement is met preferentially through the oxidation of long-chain FA [3,6,7,30]. Cardiac metabolism is therefore tightly regulated in response to intracellular levels of FA: increased FA availability (and concomitant PPARα activation) lead to decreased use of glucose with elevated FA oxidation and esterification rates [31]. Under physiological increases in FA influx where cardiac oxidative capacity is saturated, TAG accumulation limits the availability of FA for incorporation into lipotoxic species. In contrast, in pathologies such as obesity and diabetes in which metabolic stress becomes chronic and metabolic flexibility is impaired, FA overload leads to a compensatory but insufficient increase in cardiac FA oxidation [2,11], ultimately resulting in maladaptive TAG accumulation that promotes cardiac dysfunction [2,4,5,10]. Our work shows that *Acsl1* knockdown elicits a dramatic reversal of established ventricular steatosis and the subsequent amelioration of accompanying cardiac dysfunction. Although both total and short-term *Acsl1* inactivation result in increased LV mass [17,18], partial *Acsl1* abrogation did not exacerbate the hypertrophy observed in obese hearts. Obese hearts exhibited only mild cardiac fibrosis, in agreement with the discordant findings of other studies [32,33], and partial ACSL1 deficiency was associated with a moderate and transient upregulation in fibrosis [19] in both lean and obese mice that disappeared 5 weeks after knockout induction. Finally, partial *Acsl1* ablation drastically reduced ventricular TAG content, ameliorated cardiac dysfunction, and was protective in *ob/ob* hearts subjected to stress. Thus, the detrimental effects of chronic *Acsl1* deficiency were circumvented, revealing *Acsl1* modulation as a potential therapeutic strategy to mobilize intraventricular TAG in obesity-related cardiomyopathy.

That increased myocardial TAG depots correlate with LV hypertrophy, impaired diastolic function, and higher oxygen consumption has been established in the *ob/ob* model of obesity-associated cardiomyopathy [5,11]. However, *how* excess myocardial lipid storage is associated with heart disease is less clear, and overproduction of reactive oxygen species, mitochondrial membrane potential dissipation, impaired Ca^2+^ homeostasis, and the generation of lipotoxic and pro-apoptotic intermediates have all been postulated as likely culprits for obesity-associated heart failure [1,10,34-37]. Moreover, recent studies have focused on DAG and ceramides as the main contributors to obesity-associated cardiac dysfunction, proposing that TAG accumulation is a relatively innocuous player. Studies in murine models of cardiac-specific overexpression of ACSL1 [34] and DGAT1 [16] suggest that cardiac lipotoxicity is driven primarily by ceramides and DAG. In addition, increased myocardial ceramide levels in the presence of decreased TAG levels have been implicated in failing human myocardium, and TAG accumulation is present in patients with and without cardiac dysfunction [38-41], whereas severe heart failure can lead to reduced TAG and increased ceramides and DAG [3]. Cardiac-specific DGAT1 overexpression is thought to reduce toxicity associated with other transgenic models by reducing ceramide content without altering TAG levels [1]. Similarly, the cardiomyopathy observed in cardiac-specific ACSL1 overexpression is postulated to arise from selective loss of lipid-laden cardiomyocytes via ceramide-induced apoptosis [34]. However, pharmacological inhibition of *de novo* ceramide synthesis alleviates hypertrophy but does not restore survival in a model of cardiac-specific overexpression of lipoprotein lipase [42]. In addition, FA-induced apoptosis is triggered by the generation of ROS independently of ceramide synthesis [43], further suggesting that other mechanisms underlie cardiac pathologies associated with intramyocardial lipid accumulation. Thus, whether TAG plays an independent role in the development of cardiac dysfunction remains to be determined. Hearts from *ob/ob* mice have elevated levels of ceramides compared to lean controls, whereas DAG content is not affected [44]. In our study, a 60% reduction in ventricular ACSL activity was sufficient to reverse ventricular TAG accumulation in *ob/ob* hearts. Ceramide synthesis requires ACSL1-mediated activation of palmitate. Thus, partial cardiac-specific ACSL1 deficiency led to the mobilization of endogenous TAG stores while simultaneously blocking the availability of the newly-released FA for oxidation and/or incorporation into lipotoxic and pro-apoptotic species. The hydrolysis of existing TAG stores in *ob/ob* hearts that occurs after *Acsl1* inactivation would therefore increase cardiac DAG (which cannot be re-esterified due to the block in FA activation) in the context of already elevated ceramide levels, and should therefore further promote deterioration of cardiac function: yet function is improved. These findings suggest that TAG may indeed play a principal role in the development of obesity-associated cardiomyopathy.

Decreased calcium handling in *db/db* mice leads to reduced cardiac contractility [45]. Palmitate exposure results in mitochondrial membrane potential dissipation, increased ROS production, and impaired calcium signaling exclusively in wild type cardiomyocytes compared to *ob/ob* hearts [37], suggesting that prolonged exposure to excess FA can result in cardiomyocyte adaptation. Nonetheless, *ob/ob* mice exhibit decreased expression of sarco/endoplasmic reticulum Ca^2+^-ATPase and perturbed intracellular calcium homeostasis [4,46]. Could obesity-associated cardiac insufficiency be therefore compounded by a physical hindrance resulting from massive intramyocardial TAG accumulation? Mitochondria-associated membranes (MAMs) are sites of calcium exchange and have been implicated in the pathogenesis of cardiac hypertrophy and dysfunction [47]. Moreover, obesity-mediated intracellular lipid accumulation can elicit MAM reorganization [48], leading to tissue-specific effects. Thus, excessive ventricular TAG accumulation could lead to mechanical disruption of MAM interactions, or conversely, increased ER-mitochondria contacts, ultimately resulting in calcium signaling disruption and reduced cardiac contractility. Based on our findings, we speculate that the independent role of TAG accumulation on obesity-associated cardiomyopathy has been understated, and warrants further investigation.

## Abbreviations

ACSL: long-chain acyl-CoA synthetase;
DAG: diacylglycerol;
FA: fatty acid;
mTOR: mechanistic target of rapamycin;
mTORC1: mTOR complex-1;
MPI: mean performance index;
S6K: p70 S6 kinase;
TAG: triacylglycerol;
4E-BP1: eukaryotic translation initiation factor 4E-binding protein 1.

## Acknowledgements

No persons besides the authors have made substantial contributions to this manuscript. We acknowledge Brian Cooley and the UNC Animal Surgery Core Laboratory for echocardiography and Doppler analyses and helpful discussions throughout the course of this work.

## Sources of Funding

This work was supported by NIH grants DK59935 (RAC), the UNC Nutrition Obesity Research Center DK056350, and 12GRNT12030144 (RAC) from the American Heart Association Mid-Atlantic Division.

## Disclosures

Florencia Pascual, Trisha J. Grevengoed, Liyang Zhao, Monte S. Willis, and Rosalind A. Coleman: none.

